# LIONS: Analysis Suite for Detecting and Quantifying Transposable Element Initiated Transcription from RNA-seq

**DOI:** 10.1101/149864

**Authors:** Artem Babaian, Richard Thompson, Jake Lever, Liane Gagnier, Mohammad M. Karimi, Dixie L. Mager

## Abstract

**Summary:** Transposable Elements (TEs) influence the evolution of novel transcriptional networks yet the specific and meaningful interpretation of how TE-initiation events contribute to the transcriptome has been marred by computational and methodological deficiencies. We developed *LIONS* for the analysis of paired-end RNA-seq data to specifically detect and quantify TE-initiated transcripts.

**Availability:** Source code, container, test data and instruction manual are freely available at www.github.com/ababaian/LIONS.

**Supplementary information:** Supplementary data are available at Bioinformatics online.

## 1 INTRODUCTION

A major fraction of the human genome is composed of transposable elements (TEs), which contain promoters, enhancers and other cis-regulatory sequences. TEs can be viewed as a dispersed reservoir of regulatory sequences from which transcriptional innovation arises (Chuong et al., 2017; Rebollo et al., 2012).

TEs near genes may gain regulatory function over evolutionary time as alternative promoters. Interestingly, during cancer evolution, normally dormant TE promoters can be exploited to express a protooncogene. Such “onco-exaptations” have been identified for the expression of *CSF1R* and *IRF5* in Hodgkin Lymphoma (HL), *FABP7* in Diffuse Large B-cell Lymphoma, *IL-33* in Colorectal cancer, and *ALK* in melanoma, among others (reviewed in Babaian and Mager, 2016).

Previous transcriptome-wide studies designed to detect TE-derived promoters have analysed annotated mRNAs (van de Lagemaat et al., 2003), ESTs (Nigumann et al., 2002), assembled transcripts (Huda and Bushel, 2013; Kapusta et al., 2013; Kelley and Rinn, 2012), short Cap Analysis Gene Expression (CAGE) tags (Faulkner et al., 2009), paired-end ditag sequences (Conley et al., 2008), ‘chimeric-fragment’ RNA-seq screening (Karimi et al., 2011; Wang et al., 2016) or targeted TE initiation events such as ERV9- (Sokol et al., 2015) or L1-driven transcription (Cruickshanks and Tufarelli, 2009).

While these methods have proved insightful, they each have limitations. Notably, 5′ CAGE is the clearest measure of transcription start sites (TSSs) but provides insufficient information on the resultant transcript structure. RNA-seq assembly methods may not identify the true 5’ end of transcripts or suffer from a high false positive rate due to TE exonization events. Moreover, none of the aforementioned studies quantify the relative contribution of the TE-initiated isoforms to overall transcript expression when alternative promoters exist. Therefore, effective TE-initiating transcript screens have required extensive human-inspection and have failed to provide a quantitative assessment of TEs initiating transcription.

To quantitatively measure and compare the contribution of TE promoters to normal and cancer transcriptomes, we developed *LIONS* which uses paired-end RNA-seq data to rapidly measure contributions of TE-initiated transcripts. *LIONS* generates a set of TE-initiated transcripts in each library, and measures the commonalities and differences between cancer and normal groups.

## 2 METHODS

Aligned or unaligned RNA-seq data along with a reference genome, gene and repeat annotations are inputs for the classification of TE-initiated transcripts (Figure 1A). For each RNA-seq library, a standard (.lion) file of TE-initiated transcripts output, which can then be biologically grouped for comparison (i.e. cancer vs. normal) (Figure 1B). *LIONS* consists of the following three modules; data initialization; classification of TE-initiation events; and comparison between biological groups (Figure 1).

**Figure 1:**
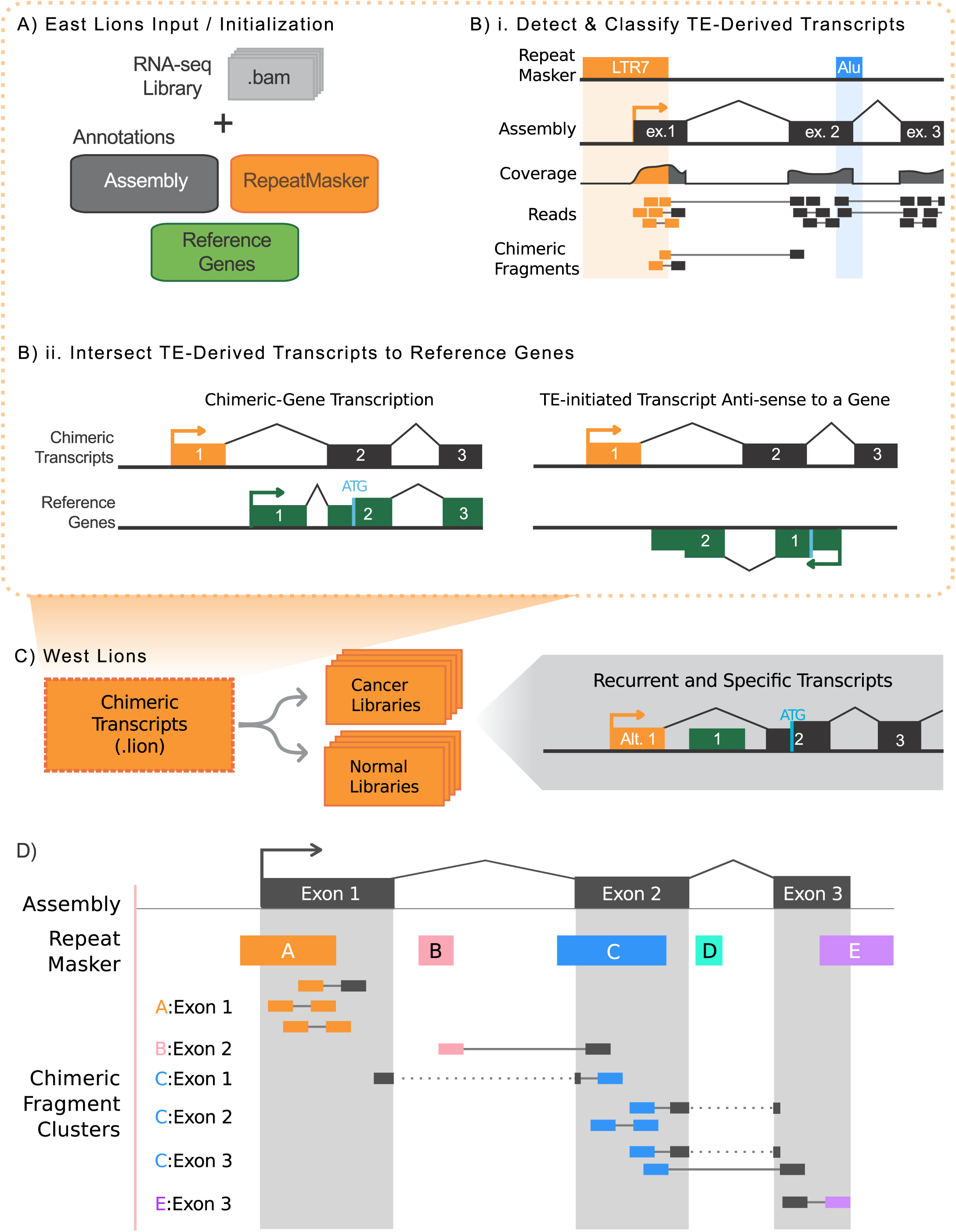
The LIONS workflow in three modules; A) ‘East Lion’ data initialization, alignment and assembly; B) Classification of TE-derived transcripts and cross-reference to protein coding gene set; and C) ‘West Lion’ analysis of biological grouping of TE-initiation sets. D) Chimeric fragment clusters consistent with transcriptional initiation (orange) are enriched and those with passive exonization (blue) or termination (purple) are depleted. Instances in which a TE-initiates a minor isoform (pink) can be included or excluded based on the ‘TE-contribution’ parameter.

### A. Initialization, Alignment, and Assembly

For an accurate measurement of TE-initiated transcripts from RNA-seq, the East Lion module was developed (Figure 1). Input is a set of paired-end RNA sequencing data in fastq or bam format, a reference genome, a RepeatMasker table (http://www.repeatmasker.org/), and a reference coding gene set. The datasets can be biologically or technically grouped or analysed individually. Optional alignment/assembly is performed with parameter-optimized *tophat2* suite (Kim et al., 2013) and secondary alignments for multi-mapping reads are retained and flagged. On systems supporting *qsub* parallelization, each library is aligned in parallel with multiple threading allowing for rapid analysis of large datasets. An optional *ab* initio assembly is constructed to reduce false-positive TE-initiation calls (relative to using a reference gene set only). The alignment and assembly is then processed to generate basic statistics for each exon and TE, such as read-coverage and RPKM expression for the repeat and genic exons which are used for transcript classification.

### B. Chimeric Fragment Cluster Analysis

To search the sequencing data for potential TE-exon interactions, each TE-exon pair for which a chimeric fragment cluster exists is considered. Briefly, a chimeric fragment cluster is a set of paired-end reads in which one read maps to a TE and its pair maps to an exon from the assembly (Figure 1D). These TE-exon pairs form the basis for classification into one of three cases; TE-initiation, TE-exonization or TE-termination of the transcript (Figure 1D).

Classification requires a series of values which are then processed by the classification algorithm (Supplementary Figure 1 and 2). First, the relative position of the TE and exon boundaries with respect to the direction of transcription is compared and only intersection cases which support TE-initiation are retained. A “thread ratio” is calculated, the ratio of read pairs in which one read maps outside of a TE in either the downstream or upstream direction. A high thread ratio distinguishes TE-initiations from TE-exonizations, that is to say, if a TE initiates transcription, then there should exist a strong bias towards the number of read-pairs downstream of the element. For the detection of TE-initiated transcripts of high biological significance, further restrictions are imposed. Single exon contigs are excluded (i.e. retained introns and low abundance lincRNA). To discard rare TE-initiated isoforms when a highly expressed isoform exists, TE contribution is estimated by peak-coverage within the TE divided by the peak coverage of its interacting exon. Together these values form the basis on which TE-initiation, TE-exonization or TE-termination can be resolved.

The parameters for the classification algorithm of TE-exon interactions can be user-customized (Supplementary Figure 2). The default set of parameters termed, ‘oncoexaptation’ were manually defined by extensive human inspection of the training ENCODE sequencing data and cross-referenced with ChIP-seq and CAGE data. The default parameters are selected to specifically detect high-abundance isoforms of TE-initiated transcripts with a significant contribution to overall gene expression. These are conservative but offer the most biologically relevant results with respect to cancer biology.

TE-initiated transcripts can be further sub-classified by their intersection to a set of protein-coding genes into (1) chimeric transcripts: TE-initiated transcripts which transcribe in the sense-orientation into a neighbouring protein-coding gene; (2) anti-sense TE-transcripts: non-coding TE-initiated transcripts which run anti-sense to a protein-coding gene; and (3) long intergenic non-coding TE-transcripts which don’t overlap a known protein-coding gene. Of particular interest to cancer biology are chimeric transcripts that result in the overexpression of potential oncogenes. Alternative filtering settings exist and are continually added based on the experimental demand. Two such settings are “screen”, which is sensitive but error-prone (e.g. exonizations called as initiations), and “drivers”, which detects TE-initiated transcripts that are exclusively transcribed from TEs. Each of these settings is customizable and can be tailored towards individual project requirements.

These analyses are performed independently for each RNA-seq library and a standard .lion file is created. Sets of .lion files (that is sets of RNA-seq library analyses) are then grouped into a merged .lion<u>s</u> file for set-based comparisons.

### C. Recurrent and Group-specific TE-promoters

Grouping and comparing sets of TE-initiated transcripts is of central importance to understanding the biology of their activity. TE-initiated transcripts are more variable than non-TE transcripts across biological replicates (Faulkner et al., 2009) and therefore the signals from individual transcriptomes are noisy. It is reasonable that grouping TE-initiated transcripts across biological replicates and asking which transcripts are recurrent will enrich for TE-initiated transcripts of consequence (see review (Babaian and Mager, 2016). In a similar line of reasoning, comparing one biological group against another can identify TE-initiated transcripts, or even classes of TEs, that are more transcriptionally active in one group of transcriptomes against another.

To detect recurrent TE-initiated transcripts between libraries with different assemblies, the West Lion was developed. Given the set of TEs that initiate transcription, the “recurrence” parameter is the number of libraries within a biological group that a given TE initiating transcription is required to be present. In contrast, the “specificity” parameter is the number of comparison (control) libraries the initiating TE is present in. Together, TEs which have greater than the recurrent cut-off and less than the specificity parameter cutoff are considered recurrent and specific TE-initiated transcripts for a group (Figure 1B).

## 3 EVALUATION

To evaluate the accuracy of LIONS-classified TE-initiations, a set of Hodgkin Lymphoma (HL) specific and recurrent (relative to B-cell controls) chimeric transcripts were assayed by RT-PCR (Supplementary Figure 3). HL and B-cell control cell culture, RNA isolation and cDNA synthesis was performed as previously described (Babaian et al., 2016). *In silico* predictions by *LIONS* were largely in agreement with RNA assayed by RT-PCR at 55.4% and 89.2% sensitivity and specificity respectively (Supplementary Figure 3). It is expected that the sensitivity of RT-PCR which reaches single-molecule sensitivity is substantially greater than RNA-seq. Altogether, LIONS is able to detect a highly specific set of TE-initiated transcripts from RNA-seq data. The detected set is enriched for higher expressed transcripts which, in a biological context such as cancer, are expected to be more relevant to oncogenesis and warrant further biological investigation.

## Acknowledgements

We thank Matt Lorincz and Rita Rebollo for helpful suggestions during the course of this work. We also thank Frances Lock for help with RNA preparation.

## References

Babaian, A., and Mager, D.L. (2016). Endogenous retroviral promoter exaptation in human cancer. Mob. DNA 7, 24.

Babaian, A., Romanish, M.T., Gagnier, L., Kuo, L.Y., Karimi, M.M., Steidl, C., and Mager, D.L. (2016). Onco-exaptation of an endogenous retroviral LTR drives IRF5 expression in Hodgkin lymphoma. Oncogene 35, 2542–2546.

Chuong, E.B., Elde, N.C., and Feschotte, C. (2017). Regulatory activities of transposable elements: from conflicts to benefits. Nat. Rev. Genet. 18, 71–86.

Conley, A.B., Piriyapongsa, J., and Jordan, I.K. (2008). Retroviral promoters in the human genome. Bioinforma. Oxf. Engl. 24, 1563–1567.

Cruickshanks, H.A., and Tufarelli, C. (2009). Isolation of cancer-specific chimeric transcripts induced by hypomethylation of the LINE-1 antisense promoter. Genomics 94, 397–406.

Faulkner, G.J., Kimura, Y., Daub, C.O., Wani, S., Plessy, C., Irvine, K.M., Schroder, K., Cloonan, N., Steptoe, A.L., Lassmann, T., et al. (2009). The regulated retrotransposon transcriptome of mammalian cells. Nat. Genet. 41, 563–571.

Huda, A., and Bushel, P.R. (2013). Widespread Exonization of Transposable Elements in Human Coding Sequences is Associated with Epigenetic Regulation of Transcription. Transcr. Open Access 1

Kapusta, A., Kronenberg, Z., Lynch, V.J., Zhuo, X., Ramsay, L., Bourque, G., Yandell, M., and Feschotte, C. (2013). Transposable Elements Are Major Contributors to the Origin, Diversification, and Regulation of Vertebrate Long Noncoding RNAs. PLoS Genet 9, e1003470.

Karimi, M.M., Goyal, P., Maksakova, I.A., Bilenky, M., Leung, D., Tang, J.X., Shinkai, Y., Mager, D.L., Jones, S., Hirst, M., et al. (2011). DNA methylation and SETDB1/H3K9me3 regulate predominantly distinct sets of genes, retroelements, and chimeric transcripts in mESCs. Cell Stem Cell 8, 676–687.

Kelley, D., and Rinn, J. (2012). Transposable elements reveal a stem cell-specific class of long noncoding RNAs. Genome Biol. 13, R107.

Kim, D., Pertea, G., Trapnell, C., Pimentel, H., Kelley, R., and Salzberg, S.L. (2013). TopHat2: accurate alignment of transcriptomes in the presence of insertions, deletions and gene fusions. Genome Biol. 14, R36.

van de Lagemaat, L.N., Landry, J.-R., Mager, D.L., and Medstrand, P. (2003). Transposable elements in mammals promote regulatory variation and diversification of genes with specialized functions. Trends Genet. TIG 19, 530–536.

Nigumann, P., Redik, K., Mätlik, K., and Speek, M. (2002). Many human genes are transcribed from the antisense promoter of L1 retrotransposon. Genomics 79, 628–634.

Rebollo, R., Romanish, M.T., and Mager, D.L. (2012). Transposable elements: an abundant and natural source of regulatory sequences for host genes. Annu. Rev. Genet. 46, 21–42.

Sokol, M., Jessen, K.M., and Pedersen, F.S. (2015). Human endogenous retroviruses sustain complex and cooperative regulation of gene-containing loci and unannotated megabase-sized regions. Retrovirology 12, 32.

Wang, T., Santos, J.H., Feng, J., Fargo, D.C., Shen, L., Riadi, G., Keeley, E., Rosh, Z.S., Nestler, E.J., and Woychik, R.P. (2016). A Novel Analytical Strategy to Identify Fusion Transcripts between Repetitive Elements and Protein Coding-Exons Using RNA-Seq. PLoS ONE 11, e0159028.

